# Evolutionary relationships among *Massospora* spp. (Entomophthorales), obligate pathogens of cicadas

**DOI:** 10.1101/811836

**Authors:** Angie M. Macias, David M. Geiser, Jason E. Stajich, Piotr Łukasik, Claudio Veloso, DeAnna C. Bublitz, Matthew C. Berger, Greg R. Boyce, Kathie Hodge, Matt T. Kasson

## Abstract

The fungal genus *Massospora* (Zoopagomycota: Entomophthorales) includes more than a dozen obligate, sexually transmissible pathogenic species that infect cicadas (Hemiptera) worldwide. At least two species are known to produce psychoactive compounds during infection, which has garnered considerable interest for this enigmatic genus. As with many Entomophthorales, the evolutionary relationships and host associations of *Massospora* spp. are not well understood. The acquisition of *M. diceroproctae* from Arizona, *M. tettigatis* from Chile, and *M. platypediae* from California and Colorado provided an opportunity to conduct molecular phylogenetic analyses and morphological studies to investigate if these fungi represent a monophyletic group and delimit species boundaries. In a three-locus phylogenetic analysis including the D1–D2 domains of the nuclear 28S rRNA gene (28S), *elongation factor 1 alpha-like* (EFL), and *beta-tubulin* (BTUB), *Massospora* was resolved in a strongly supported monophyletic group containing four well-supported genealogically exclusive lineages, based on two of three methods of phylogenetic inference. There was incongruence among the single-gene trees: two methods of phylogenetic inference recovered trees with either the same topology as the 3-gene concatenated tree (EFL), or a basal polytomy (28S, BTUB). *Massospora levispora* and *M. platypediae* isolates formed a single lineage in all analyses and are synonymized here as *M. levispora*. *Massospora diceroproctae* was sister to *M. cicadina* in all three single-gene trees and on an extremely long branch relative to the other *Massospora*, and even the outgroup taxa, which may reflect an accelerated rate of molecular evolution and/or incomplete taxa sampling. The results of the morphological study presented here indicate that spore measurements may not be phylogenetically or diagnostically informative. Despite recent advances in understanding the ecology of *Massospora*, much about its host range and diversity remains unexplored. The emerging phylogenetic framework can provide a foundation for exploring co-evolutionary relationships with cicada hosts and the evolution of behavior-altering compounds.

## Introduction

The Entomophthorales (Zoopagomycota) are among the most important arthropod-destroying fungi (Spatafora *et al*. 2016). Many North American Entomophthorales were first described by Thaxter (1888) more than a century ago. Well-known examples include *Entomophthora muscae*, causal agent of “summit disease” of numerous fly genera (Fresenius 1856, Elya *et al*. 2018) and *Entomophaga maimaiga*, a virulent pathogen and biological control agent of gypsy moth (Soper *et al*. 1988, Hajek *et al*. 1990).

Due to the ephemeral nature, obligate lifestyle, and large genome size of the Entomophthorales, these fungi are grossly underrepresented in phylogenetic studies (Spatafora *et al*. 2016, Gryganskyi *et al*. 2017). Only recently have a select few been formally investigated using molecular phylogenetics (Gryganskyi *et al*. 2012, 2013), including the recently described *Arthrophaga myriapodina*, a lethal summit disease pathogen of polydesmid millipedes (Hodge *et al*. 2017) and *Massospora*, an active host transmission pathogen of numerous cicada species (Boyce *et al*. 2019). In total, the Entomophthorales includes some 12 accepted genera with 237 species, including *Massospora* with 13 established species (Index Fungorum and MycoBank).

*Massospora* was first described anecdotally by Dr. Joseph Leidy (1851), who noted an undescribed fungal disease of periodical cicadas in the eastern U.S.: “[*Magi*]*cicada septendecim* was subject to a fungous disease” and observed that “the posterior part of the abdomen…in several instances [was] filled with a mass of oval spore-like bodies” (Leidy 1851). *Massospora* was formally established by Peck (1879) with the description of *M. cicadina* from a periodical cicada (*Magicicada septendecim*) collected in New York, USA in 1877. Following Peck’s description, Thaxter (1888) recognized *Massospora* as a member of the Entomophthorales.

Research on *Massospora* gained momentum in the twentieth century with spore development studies (Speare 1921, Goldstein 1929) and the description of ten new species in the Western Hemisphere (Ciferri *et al*. 1957, Soper 1963, 1974), two species from Australia and Afghanistan (Soper 1981), plus undescribed *Massospora* species from *Platypleura* sp. (Kobayashi 1951) and *Meimuna* sp. (Ohbayashi *et al*. 1999) in Japan. Today, *Massospora* includes more than a dozen obligate, sexually transmissible, pathogenic species that attack at least 24 cicada species worldwide (Soper 1963, 1974, 1981, Cooley *et al*. 2018) (Table 1). Nearly all extant *Massospora* species are associated with a single cicada genus with two exceptions. *Massospora cicadettae* is reported from *Plerapsalta incipiens, Chelapsalta puer,* and *Cicadetta. spp.* (Table 1) and *M. platypediae / M. levispora,* based on existing phylogenetic data, represent a single species infecting two genera of annual cicadas, *Platypedia* sp. and *Okanagana* sp. (Boyce *et al*. 2019). Generally, specimens of *Massospora* have been identified based on the cicada host they are found on, but this method of identification has proven unreliable given the recent finding that *M. levispora* and *M. platypediae* comprise a single species that occupies a broader geographic and host range than previously reported (Boyce *et al*. 2019). Until the host associations and fungus names are confirmed with detailed molecular studies, identifications based solely on host associations should be viewed with skepticism.

**Table 1:**
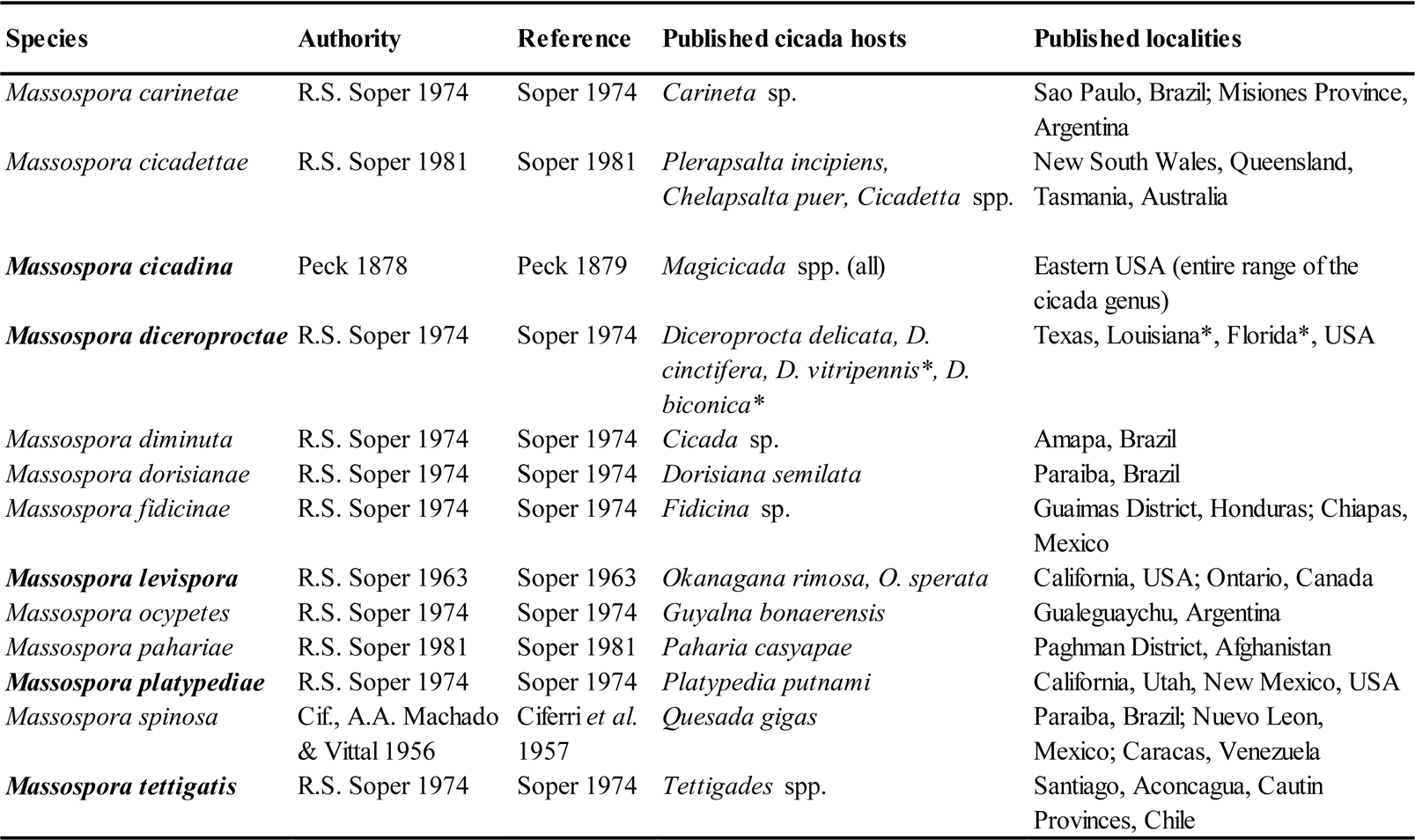
Information about the currently accepted *Massospora* species, including host information and historic collection localities. Bold font indicates the species used in this study. *: Soper is not sure if these collections represent *M. diceroproctae* or a novel *Massospora* species, as they were found far from the type locality (Soper 1974).

The life cycles of individual *Massospora* species are closely tied to the life cycle of the cicada host. Mature cicada nymphs are believed to be infected by resting spores encountered underground during construction of their vertical emergence burrows (Soper *et al*. 1976a). These nymphs then emerge, eclose into adults, and over a period of days develop infections in their abdomen. These infections become more conspicuous as the fungus destroys the cicada’s abdominal intersegmental membranes, inciting a progressive sloughing off of sclerites that reveals a large fungal mass (Figure 1). Conidia are passively disseminated during mating attempts or flights, or possibly in crowded settings where high densities of cicadas promote close contact (Soper 1963, Cooley *et al*. 2018). Cicadas infected by conidia develop secondary infections (Soper *et al*. 1976b, Cooley *et al*. 2018), resulting in the production of resting spores inside cicada hosts. These resting spores are incorporated back into the soil to infect new cohorts of cicadas as they emerge in later years (Soper *et al*. 1976a).

**Figure 1:**
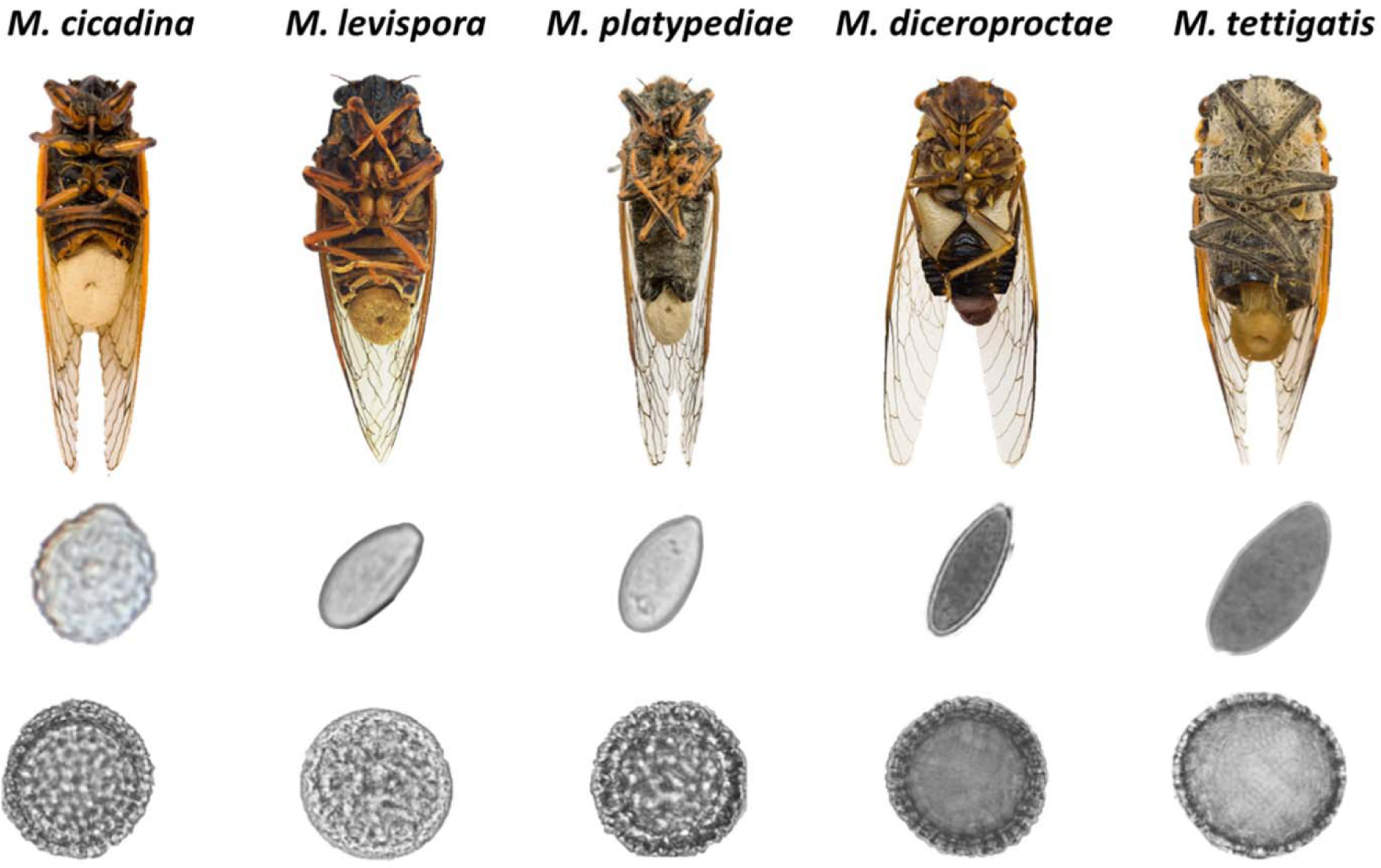
Photographs of cicada hosts (top), conidia (middle), and resting spores (bottom) of the *Massospora* species used in this study. Cicada hosts, from left to right: *Magicicada septendecim*, *Okanagana rimosa*, *Platypedia putnami*, *Diceroprocta semicincta*, *Tettigades* sp. Images are not to scale.

Complex infection and transmission strategies that involve manipulation of host behavior are notable in the Entomophthorales, including several cases of summiting behaviors and active host transmission (Roy *et al*. 2006, Hughes *et al*. 2016, Gryganskyi *et al*. 2017, Hodge *et al*. 2017, Boyce *et al*. 2019). The rarer of these two transmission behaviors, active host transmission (AHT), involves infected living hosts that directly transmit spores to new hosts (Roy *et al*. 2006). AHT behavior in *Massospora* is thought to be chemically induced (Boyce *et al*. 2019), and includes hypersexual behavior where infected male cicadas mimic female-specific behaviors to attract copulation attempts from other males (Cooley *et al*. 2018). *Massospora* and *Strongwellsea,* a fly pathogen, are the only two genera where all species are known to induce AHT behavior in their hosts, although AHT has also been reported in select species of *Entomophthora* (*E. erupta* and *E. thripidum*) and *Entomophaga* (*E. kansana*) (Roy *et al*. 2006). However, the identity and phylogenetic placement of these latter species have not been molecularly resolved (Gryganskyi *et al*. 2012, 2013). Given this taxonomic uncertainty coupled with the occurrence of both AHT and summit disease in *Entomophthora* and *Entomophaga,* the evolutionary history of AHT among members of the Entomophthoraceae should be further investigated (Boyce *et al*. 2019). More specifically, is AHT the ancestral state for the Entomophthoraceae or has it evolved several times among *Massospora, Strongwellsea, Entomophthora,* and *Entomophaga*?

Multi-locus phylogenetics using few loci can serve as a rapid, cost-effective screening tool to inform further research using genomic, transcriptomic and metabolomic approaches.

Ultimately, genomics-based approaches offer superior phylogenetic resolution, but Entomophthorales genomes are difficult to obtain for several reasons. Compared to other fungi, some Entomophthorales genomes are massive in size, including the publicly available *Entomophthora muscae* genome (600 Mb for NCBI: PRJNA479887) and *Zoophthora radicans* genome (655 Mb for JGI: ATCC 208865) (Nordberg *et al*. 2014, Elya *et al*. 2018). Additionally, many Entomophthorales are unculturable and therefore, impure and potentially degraded environmental samples must be used. Phylogenetic studies can also help populate NCBI sequence data repositories, which are significantly underpopulated for members of the Entomophthoraceae. In total, GenBank’s nucleotide sequence repository has 616 DNA sequences for the family, excluding genomes, representing only about 20% of described species. More than 45% of these sequences are from just three taxa: *Pandora neoaphidis*, *Entomophthora muscae* sensu lato and *Zoophthora radicans*. Additionally, 30% of the 616 sequences are nuclear rDNA ITS1-5.8S-ITS2 (ITS barcode) or partial nuclear 18S rRNA gene sequences, which are not suitable for accurate phylogenetic analyses (Tang *et al*. 2007, Schoch *et al*. 2012, Demirel 2016).

In this study, we used molecular phylogenetics and morphology to further investigate three findings reported by Boyce *et al*. (2019): 1) *Massospora* is monophyletic; 2) *M. levispora* and *M. platypediae* are not genealogically exclusive; and 3) *M. levispora* and *M. platypediae* are not distinguishable based on spore measurements.

## Materials and Methods

### Sample collection & DNA extraction

The following designations are used throughout the remainder of the methods: *M. cicadina* = *Mc*, *M. diceroproctae* = *Md*, *M. levispora* = *Ml*, *M. platypediae* = *Mp*, and *M. tettigatis* = *Mt*.

Infected cicadas were obtained from various locations and collectors (Table 2). Samples from each collector were stored differently, with some samples stored dry at room temperature, some frozen in RNAlater (Invitrogen, New York) or 70-95% ethanol, and some frozen dry immediately following collection (see ‘Sample Storage’ in Supplemental Table 1 & 2). The fungal plug on each infected cicada was sampled using a sterile scalpel, or by centrifuging a solution of loose spores into a pellet. DNA was extracted using a modified Wizard kit (Short *et al*. 2015). Samples were macerated in 1.5-mL microcentrifuge tubes (Eppendorf, Germany) with 600 μL of Nuclei Lysis Solution (Promega, Wisconsin) and incubated at 65 C for 30 min, vortexing at 15 min. After cooling briefly, 200 μL of Protein Precipitation Solution (Promega, Wisconsin) was added, and samples were vortexed vigorously for 10 s. Then, samples were centrifuged for 3 min at 17,562 *g*, and the supernatant was collected and moved to fresh 1.5-mL tubes with 600 μL of 99.9% isopropanol. Tubes containing the protein pellet were discarded.

**Table 2:**
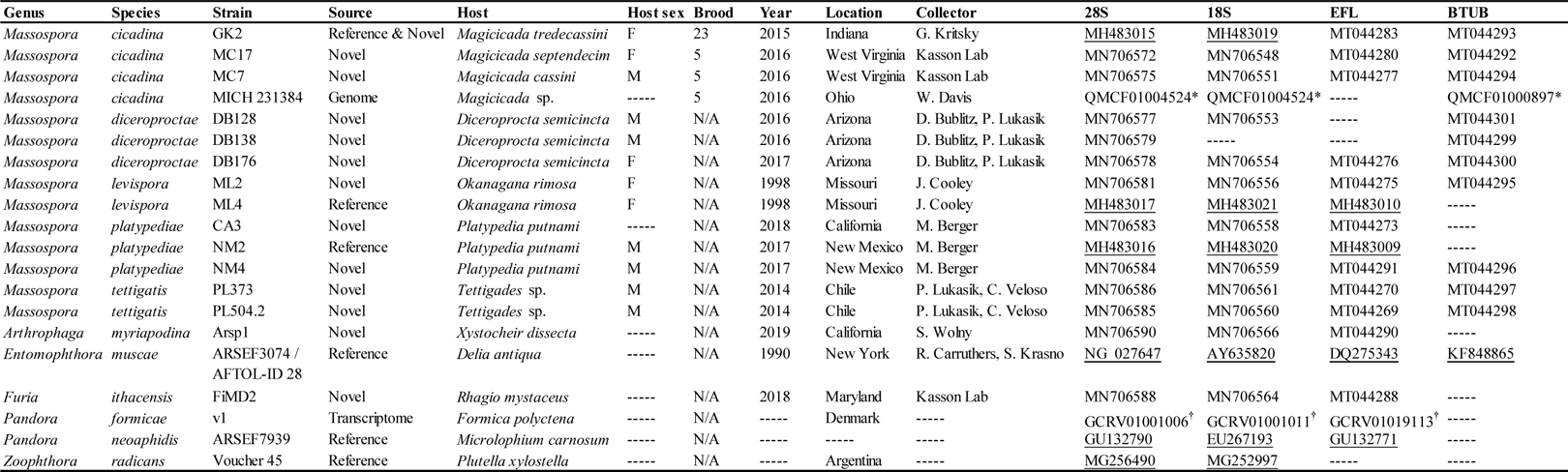
Isolates used in phylogenetic analyses and associated metadata. Reference sequences are underlined, all others are novel. * denotes NCBI whole genome shotgun sequencing project accession numbers that contain the locus of interest. † denotes NCBI Transcriptome Shotgun Assembly accession numbers that contain the locus of interest.

Sample tubes containing isopropanol were gently inverted several times and centrifuged again for 1 min at 17,562 *g.* The supernatant was discarded, leaving a DNA pellet behind. Tubes were then loaded with 600 μL of 70% ethanol and centrifuged for 1 min at 17,562 *g*. Supernatant was again discarded, and the DNA pellets were left to dry at room temperature for 20–30 min.

Finally, the DNA was resuspended in 100 μL of warmed (65 C) Elution Buffer (Alfa Aesar, Massachusetts) and stored at −20 C.

### PCR and sequencing

We targeted sequencing of the D1–D2 domains of nuclear 28S rRNA gene (28S), the V6-V9 regions of nuclear 18S rRNA gene (18S), *elongation factor 1 alpha-like* (EFL), and *beta-tubulin* (BTUB) for each sample listed in Table 2. We used existing data from GenBank for six reference strains. Additionally, six gene sequences were extracted from two assembled metagenomes from Boyce *et al*. (2019). Primer names, sequences, and full PCR protocols are listed in Supplemental Table 3. The PCR reaction volumes are as follows: 12.5 μL MyTaq^TM^ Master Mix (Bioline, United Kingdom), 10 μL molecular-grade water (G-Biosciences, Missouri), 1 μL (10 µM in IDTE, pH 8.0) each of forward and reverse primers (IDT, Iowa), and 1 μL of DNA template for a final reaction volume of 25.5 µL. PCR products were visualized via gel electrophoresis on a 1.5% w/v agarose (Amresco, Ohio) gel with 0.5% EDTA buffer (Amresco, Ohio). SYBR Gold (Invitrogen, New York) was used as the nucleic acid stain, and bands were visualized on a UV transilluminator (Bio-Rad, California). Prior to sequencing, PCR products were purified using ExoSAP-IT (Affymetrix, California): 2.2 μL of ExoSAP and 6 μL of PCR product in a 2-step reaction of 15 min at 37 C, followed by 15 min at 80 C. Purified products were Sanger sequenced (Eurofins, Alabama) with the same primers used for PCR. Sequences generated during this study are deposited in GenBank (Table 2).

### Alignments, model selection, and phylogenetic analyses

Chromatograms were quality-checked using default parameters, clipped, and manually corrected in CodonCode Aligner 5.1.5. Each gene was aligned separately using MAFFT (Katoh and Standley 2013) on the Guidance2 server (http://guidance.tau.ac.il/ver2/, Landan and Graur 2008, Sela *et al*. 2015), and individual residues with Guidance scores <0.5 were masked. An intron in 28S (positions 299–478) was deleted. Alignments are available here: http://purl.org/phylo/treebase/phylows/study/TB2:S25818

Nucleotide substitution models were chosen using AICc scores in Model Test in MEGA 7.0.16 (Kumar *et al*. 2016). Alignments of each individual gene (28S, EFL, and BTUB), and a concatenated alignment of the three genes, were used in a maximum likelihood (ML) analysis (RAxML 8.2.12, Stamatakis 2014), a maximum parsimony (MP) analysis (PAUP* 4.0a build 166, Swofford 2002), and a Bayesian inference (BI) analysis (MrBayes 3.2.5, Ronquist *et al*. 2012), for a total of 12 analyses. The default parameters of each software package were used, unless otherwise noted (see code and notes in Supplemental File 1). In brief, for ML analyses, an appropriate model was chosen, partitions were applied (for each gene in the concatenated analysis only), 1,000 bootstrap replicates were used, and the best-scoring tree was identified and bootstrapped in a single run. For MP analyses, a heuristic search with TBR swapping and 1,000 bootstrap replicates were used. For BI analyses, MrBayes was allowed to select a substitution model for each dataset, and rates were set based on results from Model Test. One cold chain and three heated chains were used for each run, and the first 25% of generations were discarded as burn-in. Each run was set for one million generations, and no additional generations were needed as the standard deviation of split frequencies fell below 0.01. Finally, runs were checked for convergence in Tracer 1.7.1 (Rambaut *et al*. 2018).

One additional tree was generated: a single-gene 18S tree using the same isolates as the 3-gene dataset, which was generated using all three methods of phylogenetic inference (see detailed methods above).

All resulting trees are available here: http://purl.org/phylo/treebase/phylows/study/TB2:S25818. Trees were viewed and prepared for publication using FigTree 1.4.4 (Rambaut 2017) and Inkscape 0.92.2 (https://www.inkscape.org/).

### Morphological study

To examine overall spore morphology, a portion of select fungal plugs (n = 63) was harvested with a sterile scalpel and mounted on a slide in lactophenol or lactophenol+cotton blue for examination with light field microscopy. Cover slips were fastened with nail polish to allow slides to be archived and re-examined when necessary. Slides were examined and photographed using a Nikon Eclipse E600 compound microscope (Nikon Instruments, New York) equipped with a Nikon Digital Sight DS-Ri1 high-resolution microscope camera. A sample of 25 spores from each slide mount were measured using Nikon NIS-Elements BR3.2 imaging software. For conidial samples, the lengths and widths of 25 conidia were recorded, and for resting spore samples, two perpendicular diameter measurements (including the epispore) were taken and averaged for 25 resting spores. Conidial measurements were taken from 45 isolates: *Mc* = 12, *Md* = 4, *Ml* = 8, *Mp* = 20, and *Mt* = 1. Resting spore measurements were taken from 18 isolates: *Mc* = 9, *Md* = 2, *Ml* = 1, *Mp* = 2, and *Mt* = 4. Raw spore measurements are available in Supplemental Table 1.

Spore measurement data was analyzed using packages dplyr (Wickham *et al*. 2019), ggplot2 (Wickham 2016), car (Fox and Weisberg 2012), userfriendlyscience (Peters *et al*. 2018), and gplots (Warnes *et al*. 2019) in R v. 3.6.1 (R Core Team, 2019). Normality was assessed using density plots and the Shapiro-Wilkes test, and equality of variance was assessed using Levene’s Test and the Fligner-Killeen test. ANOVAs and Welch’s ANOVAs (where appropriate) were performed to check for differences in spore measurements across species, and the Tukey or Games-Howell multiple-comparisons post-hoc tests (respectively) were used to identify the significant pairwise differences. A *P-*value < 0.05 was considered significant for all analyses. Reported *P*-values are Bonferroni-corrected where appropriate. R code and summarized outputs are available in Supplemental File 1.

To examine nuclei number and position in the conidia of representative *M. levispora* and *M. platypediae* specimens, spores from archived (dried or alcohol-preserved) samples were mounted in hematoxylin for observation using a Nikon Eclipse E600 phase contrast light microscope (Nikon Instruments, New York) with “PH3” and “A” filters at 100x magnification. Specimens examined for *Ml* included ML6, ML7, and ML10 (all from Michigan) and for *Mp*, NM4 and NM6 from New Mexico, CA2 from California, and CO1 and CO11 from Colorado (Supplemental Table 1). Nuclei were discernable in five *Mp* and three *Ml* specimens; other specimens had too few conidia, were in a phase of the cell cycle where the nuclei are not distinct, and/or were not receptive to staining due to age or degradation of spores. Even for samples whose conidia were receptive to staining, only a fraction of spores (< ∼25% across all samples examined) had sufficient staining to clearly identify and count nuclei. For each slide with discernable nuclei, the number and position of nuclei were recorded for 10 conidia.

## Results

### Phylogenetics

The following designations are used throughout: *M. cicadina* = *Mc*, *M. diceroproctae* = *Md*, *M. levispora* = *Ml*, *M. platypediae* = *Mp*, and *M. tettigatis* = *Mt*.

To infer evolutionary relationships among sampled taxa, several phylogenetic analyses were performed. The three individual gene trees (28S, EFL, BTUB) as well as the concatenated 3-gene tree resolved *Massospora* as a monophyletic ingroup (Figure 2). In a separate analysis, 18S placed *Md* among the outgroup taxa, and the remainder of *Massospora* was left monophyletic (Supplemental Figure 1). In all trees, *Md* resolved as a very long branch, and we attribute its occasional displacement to be a long-branch artefact, disproportionately based on signal from the 18S locus. A visual scan of all alignments indicated that differences between *Md* and other *Massospora* were distributed across all four loci, in a somewhat patchy distribution, with no indication of insertions, deletions or alignment errors being the basis of its apparent divergence. This observation together with other indications that 18S performs poorly as a phylogenetic marker for fungi (Tang *et al*. 2007, Schoch *et al*. 2012, Demirel 2016) led us to remove 18S from the concatenated analysis (Figure 2) (For 18S results see: http://purl.org/phylo/treebase/phylows/study/TB2:S25818).

**Figure 2:**
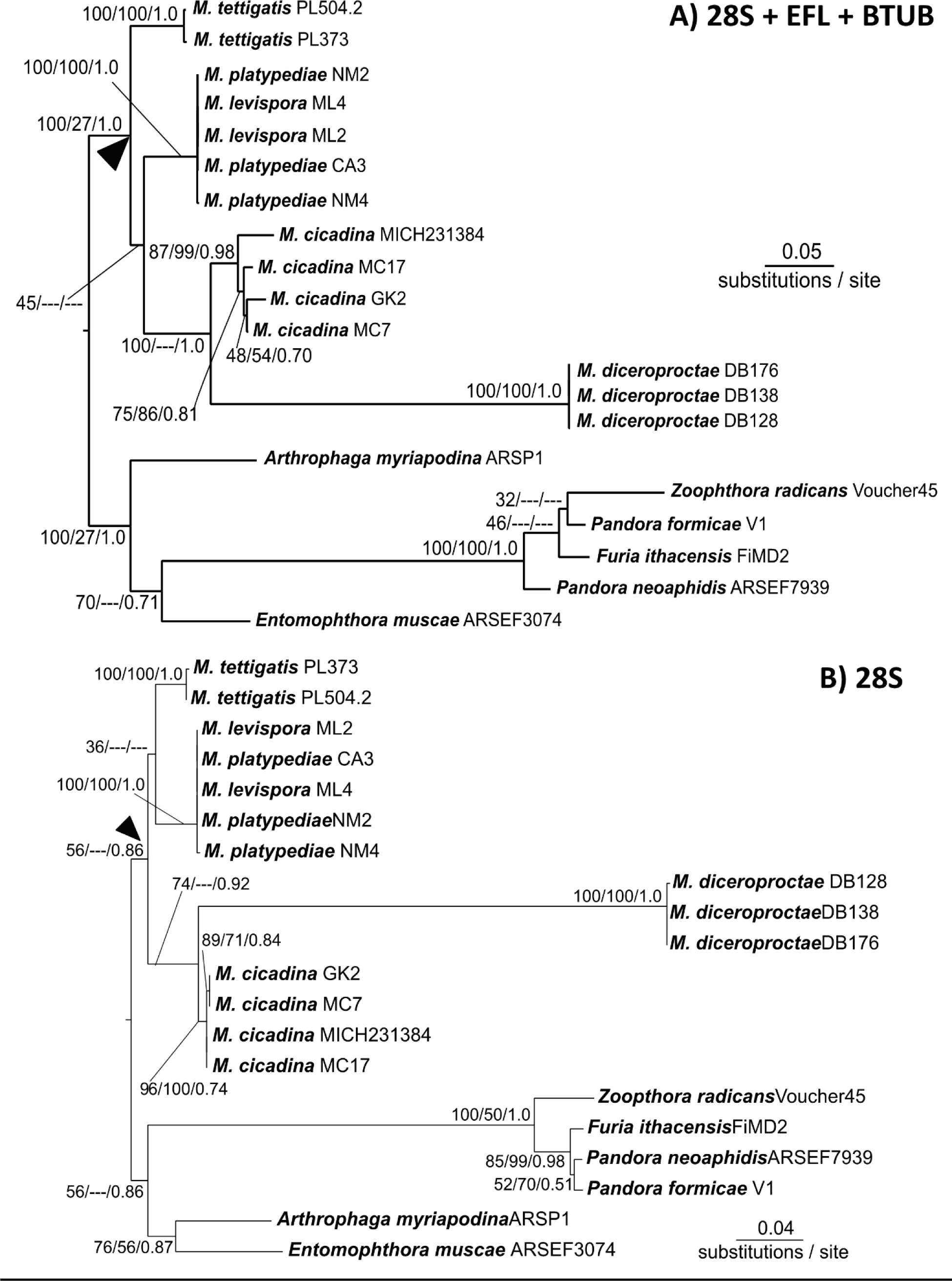
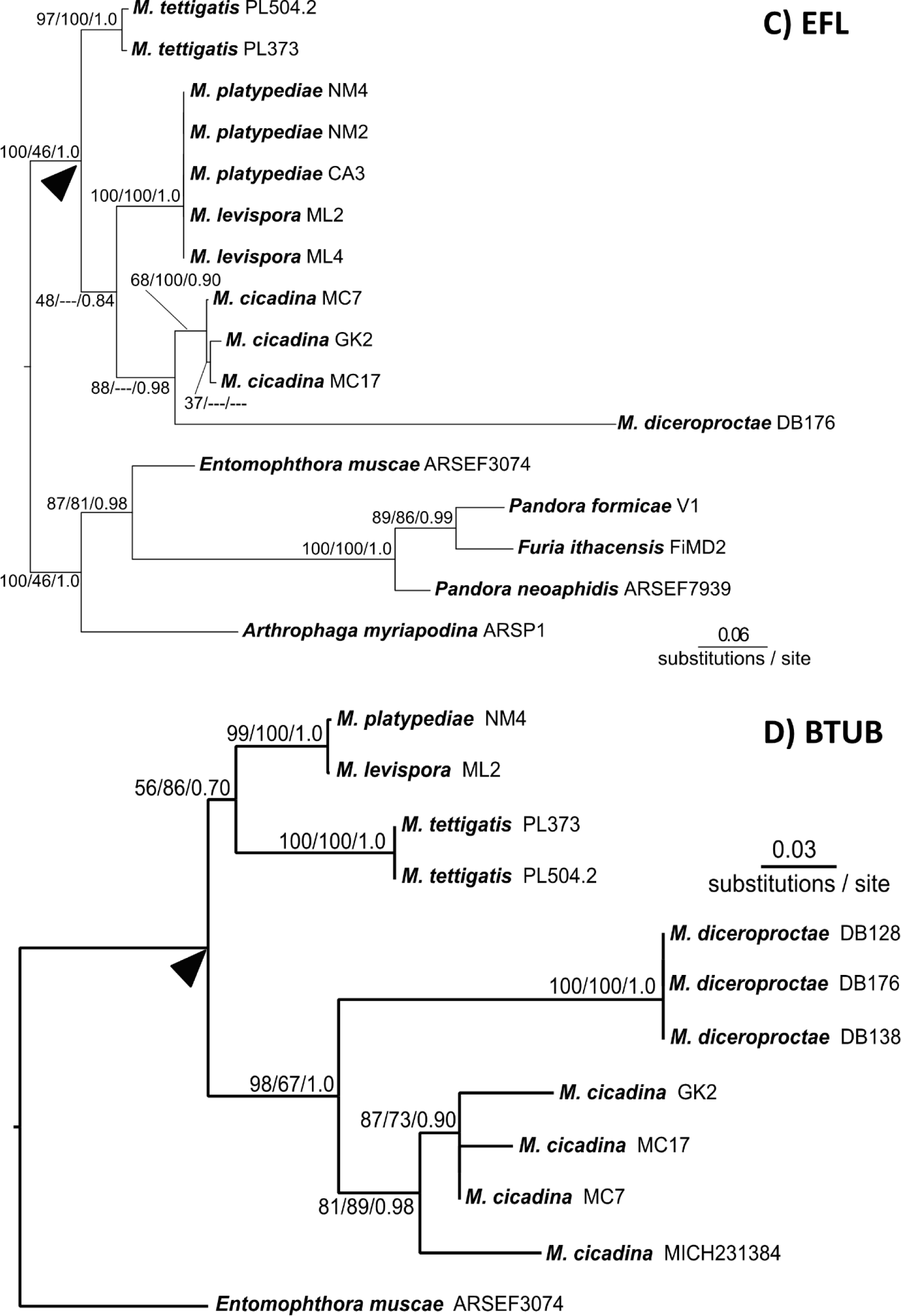
Phylogenetic trees for concatenated (A), EFL (B), 28S (C), and BTUB (D) datasets. Topology and branch lengths shown are from the ML analysis. Bootstrap support and posterior probabilities are indicated for each node supported in the ML analysis (ML/MP/BI). Dashes indicate that the node did not appear in the indicated analysis. Arrow indicates the most recent common ancestor of the genus *Massospora*.

Two of three methods of phylogenetic inference for the three-gene (28S+EFL+BTUB) concatenated dataset resolved all five *Massospora* species into a strongly supported monophyletic group (Figure 2). The third method of phylogenetic inference, MP, showed very weak bootstrap support (27%) for the genus (Supplemental Figure 1, also http://purl.org/phylo/treebase/phylows/study/TB2:S25818). A follow-up MP analysis constraining *Massospora* to be monophyletic resulted in a tree 1259 steps in length (data not shown), only four steps longer than the unconstrained analysis. Three of the *Massospora* species, *Mc*, *Md*, and *Mt*, were genealogically exclusive and had strong bootstrap support. *Massospora levispora* and *M. platypediae* did not resolve as genealogically exclusive, and instead together formed a single well-supported lineage. Within *Massospora, Mc* and *Md* formed a clade sister to the *Ml* / *Mp* lineage. *Massospora tettigatis* was recovered as the earliest diverging species of the species examined in this study.

Additional 28S sequences from specimens of *Mp* from *P. putnami* cicadas collected in 2013 in Colorado were compared using NCBI BLAST to *Mp* 28S sequences used in the 3-gene concatenated dataset. Isolates from Colorado were identical to isolates from California and New Mexico (Supplemental Table 2). Additional 28S sequences from specimens of *Mt* from three additional *Tettigades* spp. cicadas from Chile were compared using NCBI BLAST to *Mt* 28S sequences used in the 3-gene concatenated dataset. These comparisons revealed *Mt* is a single species capable of infecting diverse *Tettigates* species (Supplemental Table 2). These *Mp* and *Mt* isolates were excluded from the phylogenetic analyses due to insufficient sequence data for the other loci used.

### Morphological study

Morphological studies were conducted to permit comparisons between isolates used in this study and previously reported measurements (Soper 1963, 1974, 1981) as well as among species. Conidia and resting spore measurements were acquired from *Mp*-infected wing-banger cicadas (*Platypedia putnami*) from California and Colorado, and from *Mt*-infected *Tettigades* cicadas from Chile and *Md*-infected *Diceroprocta* cicadas from Arizona. Raw spore measurements for *Mc*, *Mp*, and *Ml* previously reported by Boyce *et al*. (2019) were also included in this study.

Conidial measurements are summarized in Figure 3 with raw spore measurements available in Supplemental Table 1. Mean, standard deviation, minimum, and maximum values for each species are reported in Supplemental Table 4. Each value is rounded to the nearest 0.5 µm. Conidial length measurements are presented as follows: Mean conidial length ± standard deviation for each species. *Mc* = 16.5 µm ± 2.0 µm, *Md* = 14.5 µm ± 2.0 µm, *Ml* = 14.5 µm ± 2.0 µm, *Mp* = 12.5 µm ± 2.0 µm, and *Mt* = 16.0 µm ± 2.0 µm. Conidial widths are reported in the same format as above, and are as follows: *Mc* = 15.0 µm ± 1.5 µm, *Md* = 7.0 µm ± 1.0 µm, *Ml* = 9.0 µm ± 1.0 µm, *Mp* = 8.0 µm ± 1.0 µm, and *Mt* = 11.5 µm ± 1.0 µm. Comparisons of mean conidial length and width among species and their statistical significance are shown in Figure 3. Overall, mean conidial width was significantly affected by species (*P* < 0.001, Welch’s ANOVA), and each individual pairwise comparison was also significant (all *P* < 0.01, Games-Howell post-hoc test). Mean conidial length was also significantly affected by species (p < 0.001, ANOVA), but mean lengths overlapped among several species (*Mt*-*Mc P* = 0.55, *Ml*-*Md P* = 1.00, *Mt*-*Md P* = 0.09, all others *P* < 0.01; Tukey’s post-hoc test) (Figure 3).

**Figure 3:**
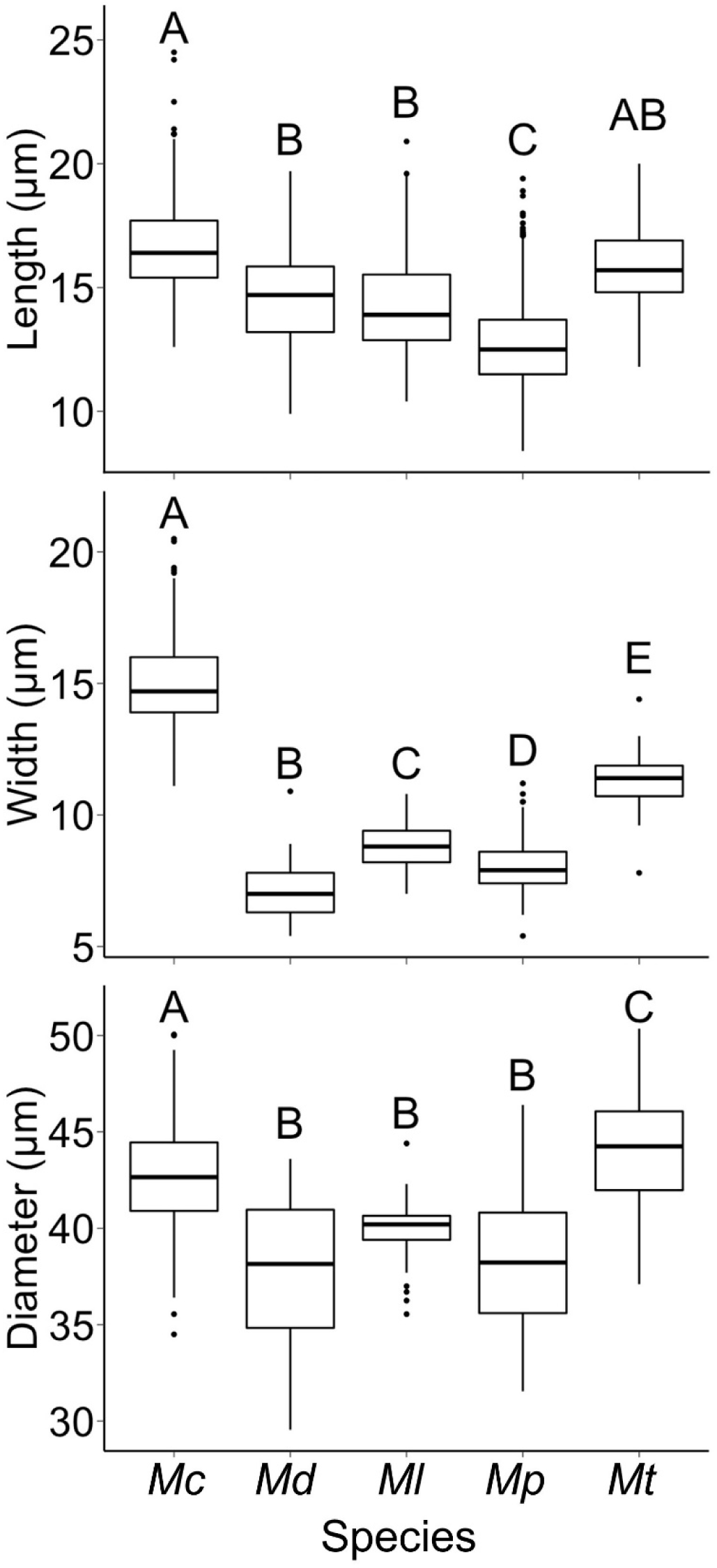
Boxplots of spore measurements used in this study. Letters indicate statistically significant differences among species. Top: Conidial length; middle: conidial width; bottom: resting spore diameter.

Unfortunately, our spore measurements cannot be statistically compared with those reported by Soper (1963, 1974, 1981) due to the fact that Soper only reported measurement means, minimums, and maximums, but not standard deviation or sample size (raw data is also unavailable). Regardless, our study found that all Soper’s mean conidial measurements fell within our reported range for each *Massospora* species (Supplemental Table 4), but not always within one standard deviation of our mean: not for *Md* conidia length, *Mp* conidia length and width, or *Mt* conidial length and width.

Resting spore measurements are summarized in Figure 3 with raw measurements available in Supplemental Table 1. Mean, standard deviation, minimum and maximum values for each species are reported in Supplemental Table 4. Resting spore diameter is reported in the same format as above, and are as follows: *Mc* = 42.5 µm ± 2.5 µm, *Md* = 38.0 µm ± 3.5 µm, *Ml* = 40.0 µm ± 2.0 µm, *Mp* = 38.5 µm ± 3.5 µm, and *Mt* = 44.0 µm ± 3.0 µm. Overall, mean resting spore diameter was significantly affected by species (*P* < 0.001, Welch’s ANOVA), but mean resting spore diameters overlapped among several species (*Ml*-*Md P* = 0.28, *Mp*-*Md P* = 1.00, *Mp*-*Ml P* = 0.44, all others *P* ≤ 0.02; Games-Howell post-hoc test) (Figure 3). Relative to Soper’s measurements, mean resting spore diameters fell within our reported range for *Mc* but not *Ml* and *Mt*, and not always within one standard deviation of our mean: not for *Ml* resting spore diameter or *Mt* resting spore diameter. Soper did not observe a resting spore stage for *Md* and *Mp* (Supplemental Table 4).

In addition to spore measurements, conidial plug color varied among species: *Md* plugs from specimens were violet to purple in color, compared to creamy white to brown plugs from all other species included in this study (Figure 1).

### Taxonomy

*Massospora levispora* and *M. platypediae* formed an unresolved clade in phylogenetic reconstructions based on 18S, 28S, and EFL, as well as the combined 4-gene tree and previous work (Boyce *et al*. 2019), suggesting these names should be considered synonyms. The two species were described from different hosts and different geographical areas: *Massospora levispora* was described from *Okanagana rimosa* cicadas collected in Ontario, Canada (Soper 1963), whereas *M. platypediae* was described from *Platypedia putnami* cicadas collected in California, New Mexico, and Utah (1974). Hosts have often been considered important in species delimitation in *Massospora*, but host specificity has seldom been experimentally studied. Morphologically, Soper’s studies determined that *Mp* had uniform broadly ellipsoidal conidia with two bipolar nuclei, whereas *Ml* had less-uniform ellipsoidal to ovoid conidia with 1-3 randomly distributed nuclei (Supplemental Table 5). No samples of *Mp* resting spore material were available at that time, but *Ml* resting spores were described as round, broadly and irregularly reticulate, and bearing many small rounded papillae discernible in scanning electron micrographs (SEM) (Soper 1974) but not in light micrographs (Soper 1963) (Supplemental Table 5).

We observed that conidial dimensions for *M. levispora* and *M. platypediae* were significantly different (Figure 3, Supplemental Table 4 & 5). Our observations confirmed the presence of ellipsoidal conidia in both species, but no ovoid conidia were observed in either species (Figure 4). For both *Ml* and *Mp,* most spore contained two medial nuclei (Supplemental Table 5). Bipolar large oil droplets were observed in some spores of both *Ml* and *Mp*. We observed for the first time the resting spores of *M. platypediae*. The spores were round with a finely reticulated rough epispore (Figure 4). We could not determine if papillae were present, due to the limitations of light microscopy. Comparing *Ml* and *Mp* resting spores, we found no significant difference in size (Figures 3, 4, and Supplemental Tables 4 & 5).

**Figure 4:**
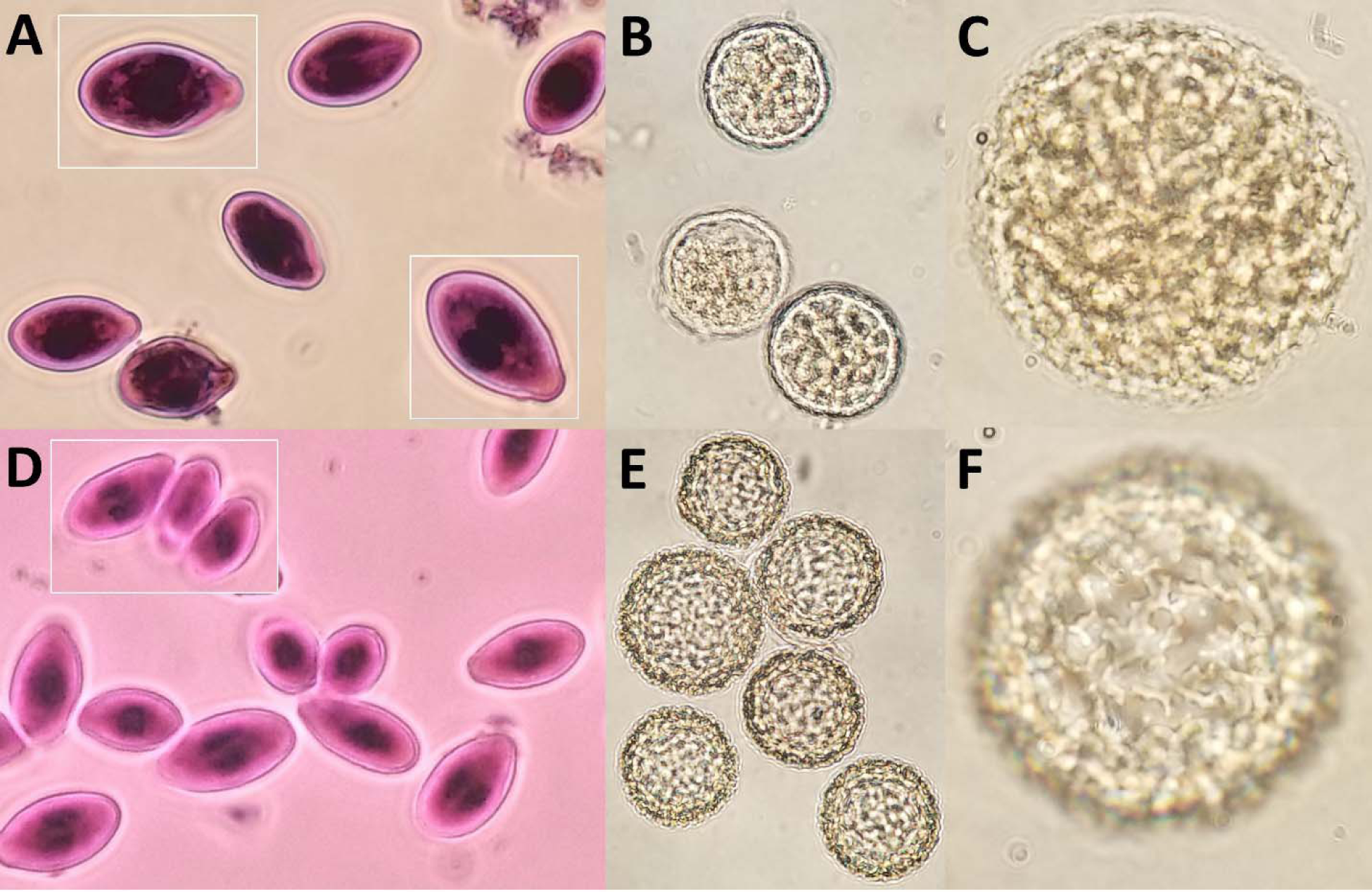
Composite of light microscopy images of *Massospora levispora* (A-C) and *M. platypediae* (D-F) conidia (A, D) and resting spores (B, C, E, F). Images shown not to scale. Conidia were mounted in hematoxylin and resting spores were mounted in lactophenol. Isolates are A) ML7, B and C) ML3, D) CO11, and E and F) NM11.

In summary, neither morphological nor phylogenetic analysis supports the recognition of two separate species, and therefore we propose the following synonymy:

***Massospora platypediae* R.S. Soper**, Mycotaxon 1 (1): 23 (1974) [MB#317412] =*Massospora levispora* R.S. Soper, Canadian Journal of Botany 41 (6): 875 (1963) [MB#333869]

## Discussion

In a recent study (Boyce *et al*. 2019), three species of *Massospora* were found to form a monophyletic group containing two genealogically exclusive lineages. In this work, we confirmed the monophyly of *Massospora*, even with the addition of two previously unavailable described *Massospora* species. At least four *Massospora* species are now well-supported according to the criteria of genealogical concordance and non-discordance (Taylor *et al*. 2000; Dettman *et al*. 2003).

The incongruence between spore morphology and molecular phylogenetics regarding the *Ml* / *Mp* lineage is intriguing. *Ml* conidia from *O. rimosa* are significantly longer (*P* < 0.01) and wider (*P* < 0.01), compared to *Mp* counterparts from *P. putnami*. Additionally, Soper’s mean conidial length and width measurements for *Mp* are not within one standard deviation of our measurements, nor are his resting spore measurements for *Ml* (Supplemental Table 4). The mountant used for spore measurements may affect spore shape and size but it is not known what mountant was used by Soper (1963, 1974, 1981). Other studies of Entomophthorales used lactophenol, aceto-orcein, or lactic acid (Humber 1976, Soper *et al*. 1988, Gryganskyi *et al*. 2013, Hodge *et al*. 2017, Małagocka *et al*. 2017). Differences among species (Fig. 3) and in comparison to Soper’s measurements (Supplemental Table 4) may also be due to differing sample ages and storage: in our study, *M. levispora* samples were stored in ethanol for 20 years whereas *M. platypediae* were stored dry and only for a few years (Boyce *et al*. 2019; Supplemental Table 1). It is not known how Soper’s samples were stored or for how long (Soper 1963, 1974, 1981). One study examining sample age and mountant used in *Strongwellsea* found that these factors have an interacting effect on spore dimensions (Humber 1976). Sample size may also be important (n = 8 for *Ml*, n = 20 for *Mp*). Previous work by Boyce *et al*. (2019) used fewer populations of *Mp* (14 isolates from one population) and found considerable overlap in both conidium and resting spore measurements for *Mp* and *Ml*, although these measurements were not statistically compared. Taken together, these studies suggest that there may be population-level variation in *Mp* spore sizes, such that sampling too few populations will result in misleading conclusions. However, this does not explain the incongruence of our phylogenetic study and morphologic study, with respect to *Mp* and *Ml*. Further sampling is needed.

*Massospora diceroproctae* was on an extremely long branch relative to the other *Massospora* species in both the 3-gene concatenated tree and the single-gene trees (Figure 2), sometimes longer than even the total branch length separating the genus *Massospora* from the most distantly related outgroups. In several parsimony-derived trees, *Md* fell among the outgroups. Some of the incongruence between MP and the other methods of phylogenetic inference observed in this study can be explained by long branch attraction (LBA) (Felsenstein 1978) acting on the *Md* clade and the outgroup clade. This result is not entirely surprising, given that MP is often more susceptible to LBA than other phylogenetic methods (O’Connor *et al*. 2010). In the 3-gene ML and BI concatenated trees (http://purl.org/phylo/treebase/phylows/study/TB2:S25818), LBA cannot explain *Mc* and *Md* forming a clade, because *Mc* is not on a long branch in this study, and did not appear on a long branch in Boyce *et al*. (2019) either, in a tree with only *Mc*, *Ml*, and *Mp*.

One possible explanation for the long branch lengths and inconsistent resolution of *Massospora* in this study is that *Md* may have experienced an accelerated rate of molecular evolution compared to all other *Massospora* species. A second, perhaps more likely explanation for long branches associated with *Md* is that the closest relatives of *Md* were not sampled here, due either to unavailability of samples, their undiscovered status, or extinction. Only five of the 12 described *Massospora* species were available for this study, and there may also be undiscovered extant taxa that would disrupt the long branches associated with *Md*. *Massospora* is not the only member of the Entomophthorales where long branches have been observed: *Batkoa* was recovered on a longer branch compared to other taxa in two separate analyses (Gryganskyi *et al*. 2012, Hodge *et al*. 2017). Similar long-branch taxa have been observed in other early diverging fungi outside the Entomophthorales, which can be partially explained by the limited taxon sampling compared to members of Basidiomycota and Ascomycota (James *et al*. 2006b, Jones *et al*. 2011).

Two *Massospora* species treated in this study, *Mt* and the *M. levispora* sensu lato have cicada hosts both belonging to the subfamily Tibicinae, whereas the hosts of *Md* and *Mc* belong to two other subfamilies, Cicadinae and Cicadettinae, respectively (Sanborn 2013, Łukasik *et al*. 2018, Marshall *et al*. 2018). Our results indicate that all three cicada subfamilies are susceptible to *Massospora*, but *Massospora* has only been molecularly confirmed from cicadas in the New World. All three subfamilies contain dozens of genera and species that have never been formally surveyed for *Massospora*. Before cophylogenetic analyses of *Massopora* and their cicada hosts can be performed to test for evidence of parallel cladogenesis, the relationships among *Massospora* species need to be better resolved through the addition of more taxa and other loci.

Given the previous findings by Boyce *et al*. (2019) that two species of *Massospora*, *Mc* and *M. levispora* sensu lato, produce psychoactive compounds during host infection, and the findings of this study that within *Massospora*, *Md*, *Mc*, and *M. levispora* sensu lato form a clade, *Md* is a likely candidate worth investigating for similar biologically active compounds.

Observations of *Md*-infected *Diceroprocta semicincta* in Arizona revealed altered calling patterns in these cicadas despite continued mating attempts (Dr. DeAnna Bublitz, personal observations). A separate personal observation of *M. diceroproctae-*infected *Diceroprocta* sp. by Dr. Jon Hastings from Big Bend National Park in Texas showed behavioral changes in infected individuals: elevated mating effort in terms of time spent signaling in males and increased likelihood to be in contact with a conspecific for males and females. Few observations exist on the behavior of *Massospora*-infected *Tettigades* cicadas, although *Mt*-infected cicadas continue mating attempts (Dr. Piotr Łukasik, personal observations). Collectively, these personal observations are intriguing, but more formal observations are needed to validate these findings.

The results of the morphological study presented here indicate that spore measurements may not be useful for species level identifications. Unfortunately, the numbers of isolates sampled for many of these species were insufficient to confidently conclude whether differences truly exist. In general, trends observed across spore measurements were incongruent with the evolutionary relationships proposed by molecular phylogenetics. For example, comparisons between *Ml* isolates and *Mp* isolates uncovered significant differences in conidium length (*P* < 0.01) and width (*P* < 0.01) (Figure 3) despite forming a single lineage based on multi-locus sequence data (Figure 2). However, resting spore diameter was not significantly different between *Ml* and *Mp* (*P* = 0.44).

In less than a decade, the research on Entomophthorales has grown significantly, leading to breakthrough discoveries on the biology and ecology of several members of this long-neglected group (Grell *et al*. 2011; Małagocka *et al*. 2015, De Fine Licht *et al*. 2017, Arnesen *et al*. 2018, Elya *et al*. 2018, Wronska *et al*. 2018, Boyce *et al*. 2019). Still, the vast majority of the Entomophthorales remain understudied. Despite recent advances in understanding the ecology of *Massospora* (Cooley *et al*. 2018, Boyce *et al*. 2019), much about the host-range and diversity of this genus is yet to be discovered. The emerging phylogenetic framework for *Massospora* provides a starting point for co-evolutionary studies with their cicada hosts and also lays a foundation for deciphering the evolution of behavior-altering compounds among *Massosopora* and close allies.

## Acknowledgements

AM was supported by The Ruby Distinguished Doctoral Fellows Program, Morgantown, WV, P.L. by National Geographic Society grant 9760-15, J.E.S. by NSF DEB 1441715, and M.T.K. by funds from West Virginia Agricultural and Forestry Experiment Station. This publication is Article No. XXXX of the West Virginia Agricultural and Forestry Experiment Station, Morgantown, West Virginia, USA, 26506.

---

**Supplemental Figure 1:**
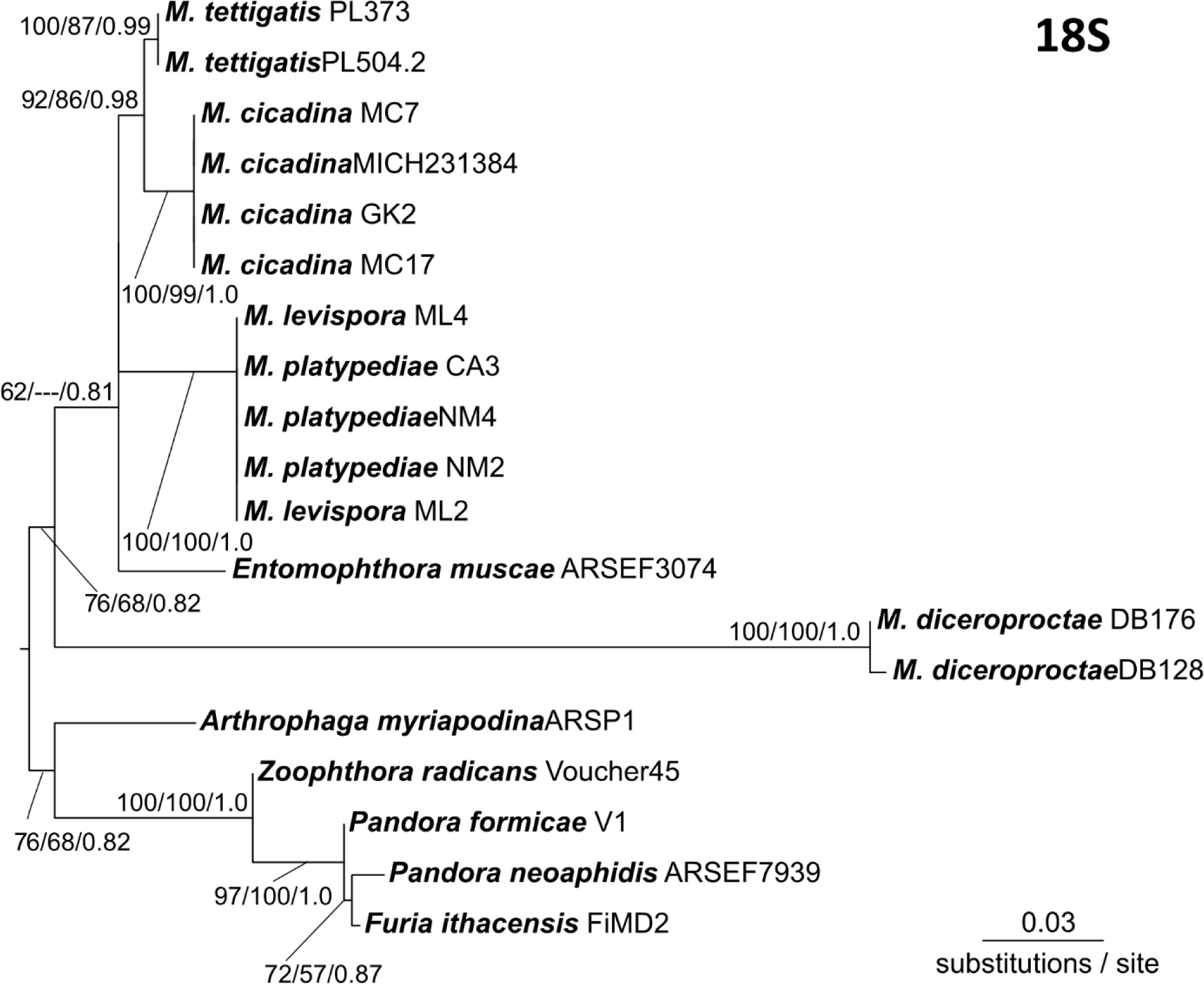
18S single-gene tree of *Massospora* spp. and outgroups. Topology and branch lengths shown are from the ML analysis. Bootstrap support and posterior probabilities are indicated for each node supported in the ML analysis (ML/MP/BI). Dashes indicate that the node did not appear in the indicated analysis.

**Supplemental File 1**: Phylogenetics code and notes for PAUP*, RAxML, and MrBayes, and R code used for spore measurement-related statistical comparisons.

